# Estimation of permutation-based metabolome-wide significance thresholds

**DOI:** 10.1101/478370

**Authors:** Alina Peluso, Robert Glen, Timothy M D Ebbels

## Abstract

**Motivation:** A key issue in the omics literature is the search for statistically significant relationships between molecular markers and phenotype. The aim is to detect disease-related discriminatory features while controlling for false positive associations at adequate power. Metabolome-wide association studies have revealed significant relationships of metabolic phenotypes with disease risk by analysing hundreds to tens of thousands of molecular variables leading to multivariate data which are highly noisy and collinear. In this context, conventional Bonferroni or Sidak multiple testing corrections are rather useful as these are valid for independent tests, while permutation procedures allow for the estimation of significance levels from the null distribution without assuming independence among features. Nevertheless, under the permutation approach the distribution of p-values may present systematic deviations from the theoretical null distribution which leads to overly conservative adjusted threshold estimates i.e. smaller than a Bonferroni or Sidak correction.

**Methods:** We make use of parametric approximation methods based on a multivariate Normal distribution to derive stable estimates of the metabolome-wide significance level. A univariate approach is applied based on a permutation procedure which effectively controls the overall type I error rate at the *α* level.

**Results:** We illustrate the approach for different model parametrizations and distributional features of the outcome measure, using both simulated and real data. We also investigate different levels of correlation within the features and between the features and the outcome.

**Availability:** MWSL is an open-source R software package for the empirical estimation of the metabolome-wide significance level available at https://github.com/AlinaPeluso/MWSL.

## 1 Introduction

One important goal in omics studies is the detection of statistically significant relationships between molecular concentrations (features) and disease phenotypes. The aim is to gauge each variable’s strength of association with disease while minimising the risk of false positive associations at adequate power. In such studies many hundreds to tens of thousands of molecular variables are collected for each individual, leading to high-dimensional multivariate data which are highly collinear. Therefore, effective methods to adjust for multiple testing are a central topic, especially in the context of Metabolome-Wide Association Studies (MWAS) (Holmes, 2008).

The aim is to adjust the observed significance level or p-value from the statistical test of each feature which is employed to detect the association between metabolic variables and disease status. In this paper we focus on the estimation of a permutation-based threshold required to control the family-wise error rate (FWER) as previously described in similar contexts (Hoggart *et al*., 2008; Chadeau-Hyam *et al*., 2010; Castagné *et al*., 2017). Although permutation procedures can yield the correct FWER even when the tests are dependent, it may happen that the permutation-based estimated threshold would be lower than those estimated with more traditional methods such as the Bonferroni or Sidak adjustment. These are known to be valid in the case of independent tests but they tend to be overly conservative in association studies in which the tests are correlated. A valid adjustment for multiple testing must account for the dependence between variables, and guarantee that the metabolome-wide significance level (MWSL) estimate would be larger than that estimated from a Bonferroni or Sidak correction. However, this is not always the case with the procedures outlined in Castagné *et al*. (2017).

To address this, we make use of approximation methods within a permutation procedure to derive a stable estimate of the MWSL, while retaining the data structure up to the 2nd order moments. This guarantees that the complex correlation structure existing in the metabolic profiling is accounted for. We illustrate the results for different model parametrizations and diverse distributional properties of the outcome, as well as by investigating different correlation levels of the features and between the features and the phenotype in real data and simulated studies. Moreover, we show that the method effectively controls the expected overall type I error rate at the *α* level.

## 2 Methods

### 2.1 Permutation-based MWSL estimation

Suppose the data consists of *n* observations, and let *Y* be the response variable, *X* = (*X*_1_,…, *X*_*M*_)^*T*^ the vector of *M* predictors or features, and *Z* = (*Z*_1_,…, *Z*_*P*_)^*T*^ the vector of *P* fixed effects covariates. The permutation-based MWSL estimation can be described as follows.

- Step (1): Shuffle i.e. re-sample without replacement, the outcome variable *Y* together with the set of fixed effects confounders *Z* if any. In this way, the *n* subjects are re-sampled under the null hypothesis of no association.
- Step (2): To estimate the relationship between the outcome and the set of features while accounting for possible confounding effects, compute *M* regression models in a univariate approach, that is by using one feature at a time. From each model store the p-value associated with the feature of interest.
- Step (3): Extract the minimum of the set of *M* p-values as this indicates the highest threshold value which would reject all *M* null hypotheses.
- Step (4): Repeat Step (1)-(3) for *K* = 10, 000 times, or at least *K* = *n/*2 times (Paparoditis *et al*., 1999). The *K* minimum p-values are the elements of the new vector *q*.
- Step (5): Sort the elements of *q*, and take the (*αK*)-value of this vector. This value is the MWSL estimate. An approximate confidence interval can be obtained by treating the true position of the MWSL estimate as a Binomial random variable with parameters *K* and *α*. Then, using the Normal approximation to the Binomial, we obtain the (1-*α*)% confidence limits by extracting the elements of *q* in positions 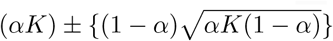.
- Step (6): Compute the effective number of tests (ENT) defined as the number of independent tests that would be required to obtain the same significance level using the Bonferroni ENT 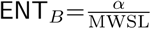, or the Sidak ENT 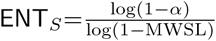 correction. The ENT estimate measures the extent that the *M* markers are non-redundant. Therefore, the ratio 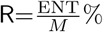 of the effective and the actual number of tests (ANT or *M*) is a measure of the dependence among features, which it is expected to be closer to 0% when highly correlated features are considered.

Similar versions of this procedure have been previously applied in different studies e.g. by Hoggart *et al*. (2008) to approximate the genome-wide significance threshold for dense SNP and resequencing data, or by Chadeau-Hyam *et al*. (2010) for urinary metabolic profiles. Recently in the context of NMR metabolic profiling studies Castagné *et al*. (2017) employed the permutation algorithm to perform a series of MWAS for serum levels of glucose, and they found ENT estimates greater than the actual number of tests, e.g. R ratio for glucose over 400%.

### 2.2 Parametric approximation methods

We investigate the properties of the permutation approach for significance level estimation, and we employ the multivariate Normal and the multivariate log-Normal distributions to describe, at least approximately, the set of correlated features to obtain stable estimates of the MWSL while effectively controlling the maximum overall type I error rate at the *α* level.

We assume that the data are already centred so that the means equal zero. Therefore, *X ∼ 𝒩*_*M*_ (*µ,* Σ^*\**^) is the multivariate Normal distribution approximation to the set of features where *µ* = E[*X*] = (E[*X*_1_],…, E[*X*_*M*_])^*T*^ = **0** is the *M*-dimensional mean vector of zero means, and Σ^*\**^ is the (*M × M*) shrinkage estimator of the covariance matrix as described by Schäfer *et al*. (2005), which is always positive definite, well-conditioned, more efficient and therefore preferred to the unbiased estimator Σ, or to the related maximum likelihood estimator Σ_ML_. When the simulation via the multivariate log-Normal distribution is considered the features are first transformed i.e. the absolute value of their minimum plus one unit is added to their original value.

The algorithm is applied to real-data and simulated scenarios to illustrate the results for different model parametrizations and distributional features of the outcome, as well as to investigate different correlation levels across features and between the features and the phenotype.

## 3 Results

### 3.1 Study of experimental metabolomics data

The MWAS approach was employed to investigate the association between human serum ^1^H NMR metabolic profiles and various clinical outcomes in the Multi-Ethnic Study of Atherosclerosis (MESA) (Bild *et al*., 2002). The data have been extensively described in Castagné *et al*. (2017).

Briefly, the cohort includes participants (51% females, 49% males), aged 44-84 years, (mean=63 years) from four different ethnic groups: Chinese-American, African-American, Hispanic, and Caucasian, all recruited between 2000-2002 at clinical centres in the United States and free of symptomatic cardiovascular disease at baseline. Demographic, medical history, anthropometric, and lifestyle data, as well as serum samples were collected, together with information on diabetes, and lipid and blood pressure treatment. Metabolic profiles were obtained using ^1^H NMR at 600 MHz and processed as detailed in Karaman *et al*. (2016). The outcomes of interest are glucose concentrations and the body mass index (BMI). Supplementary Table S1 presents the descriptive statistics for the clinical outcome measures in Supplementary Figure S1, while Supplementary Table S2 reports the descriptive statistics for the fixed effects covariates used in the study.

Three sets of NMR spectra are considered: (1) a standard water-suppressed one-dimensional spectrum (NOESY), and (2) a Carr-Purcell-Meiboom-Gill spectrum (CPMG), and (3) a lower resolution version of the CPMG data (BINNED). The BINNED version consists of *M* =655 features, while the NOESY and CPMG contain *M* =30,590 features. The BINNED data sample comprises of *n*=3,500 individuals, while the NOESY and CPMG data have *n*=3,867 participants. All MWSL calculations are performed for *α* = 0.05.

From the analysis of the BINNED data shown in Figure 1, when the real features are considered, there is instability in the estimation of the ENT across the different outcomes, and in particular the ENT estimate for glucose is above the ANT. When the data are approximated via a multivariate log-Normal or Normal as described in Section 2.2 the ENT estimates are stable across the different outcomes and remain bounded below the total number of features with an average ENT of 355 and an R ratio around 55%. To assess the validity of this result in terms of redundancy of the set of features we considered principal component analysis (PCA) as an alternative method for estimating the ENT (Cheverud, 2001; Nyholt, 2004; Li *et al*., 2005). The cumulative proportion of variance explained by the first 355 PCs is around 99%. This is consistent with the interpretation that there are approximately 355 uncorrelated features in the data.

**Figure 1:**
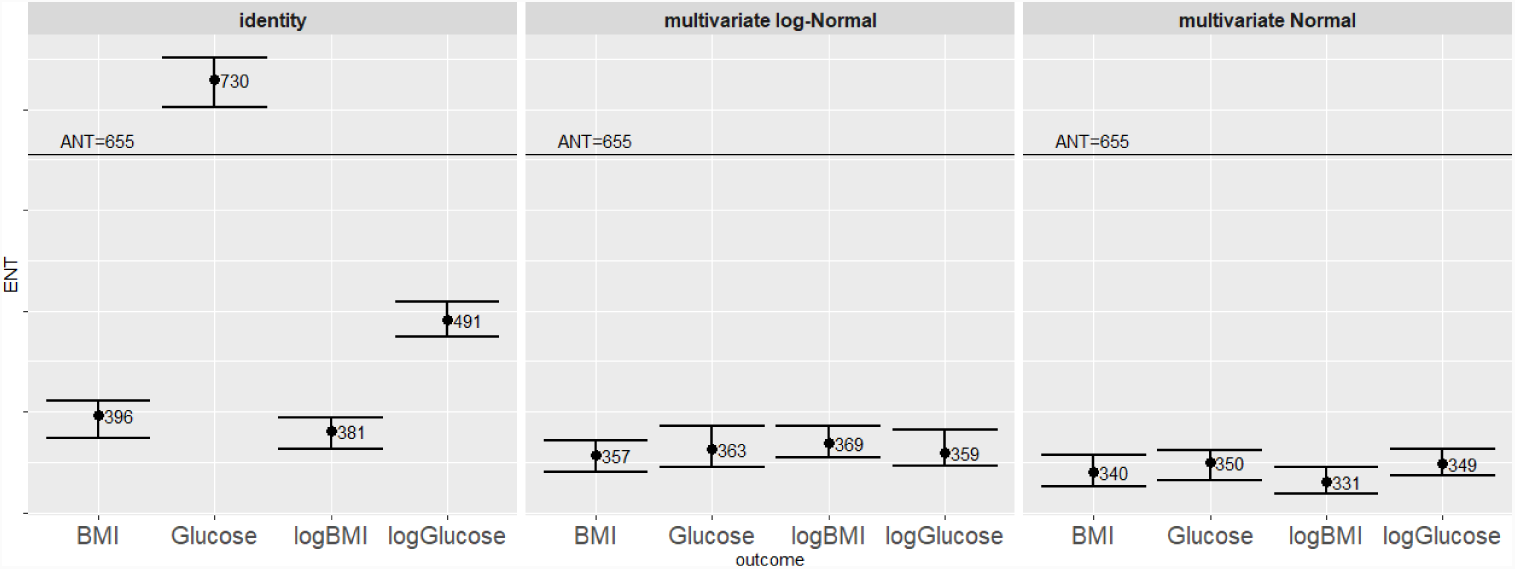
BINNED data: ENT across clinical outcome measures and for different transformations of the features: no transformation (identity), multivariate Normal, multivariate log-Normal. Error bars represent 95% confidence limits. *K*=10,000 permutations.

Figure 2 reports the ENT estimates for CPMG data. Without any transformations applied, there is a very large variation across the ENT estimates for the different outcomes, and in particular a very high and meaningless estimate for glucose levels which goes beyond R=400%. On the other hand, when the set of features is simulated from the multivariate Normal and from the multivariate log-Normal distribution the corresponding ENT estimate is below the total number of features, and stable across different outcomes with an average ENT of around 16,000 features and an R ratio around 55%. In this case the usefulness of the proposed permutation method to estimate the ENT is clear in a comparison with the PCA-based methods as proposed by Cheverud (2001), Nyholt (2004), and Li *et al*. (2005) as the ENT estimates would be constrained to the maximum number of PCs (*n* = 3, 867).

**Figure 2:**
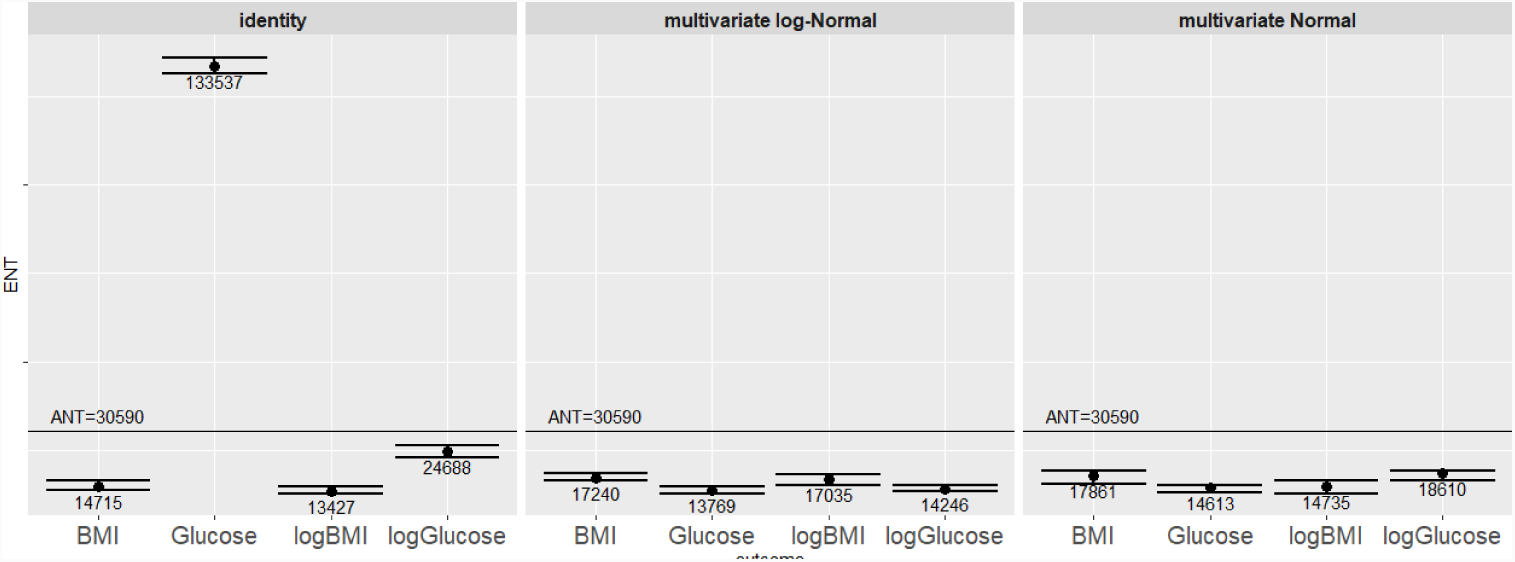
CPMG data: ENT across clinical outcome measures and for different transformations of the features: no transformation (identity), multivariate Normal, multivariate log-Normal. Error bars represent 95% confidence limits. *K*=10,000 permutations.

Figure 3 reports the ENT estimates for the NOESY data which are below R=100% but vary across outcomes when the original set of features is considered. When simulated features from the multivariate (log-) Normal distributions are considered we obtain lower ENT values than the ones from the CPMG data, with an average ENT of around 2,700 features and an R ratio around 9%. This result was expected due to the reduced influence of broad signals in CPMG spectra compared to NOESY, which is linked to a reduction in the covariance structure of the data. By applying a PCA to the NOESY data the cumulative proportion of variance explained by the first 2,700 PCs is around 99%, and this is in line with our findings.

**Figure 3:**
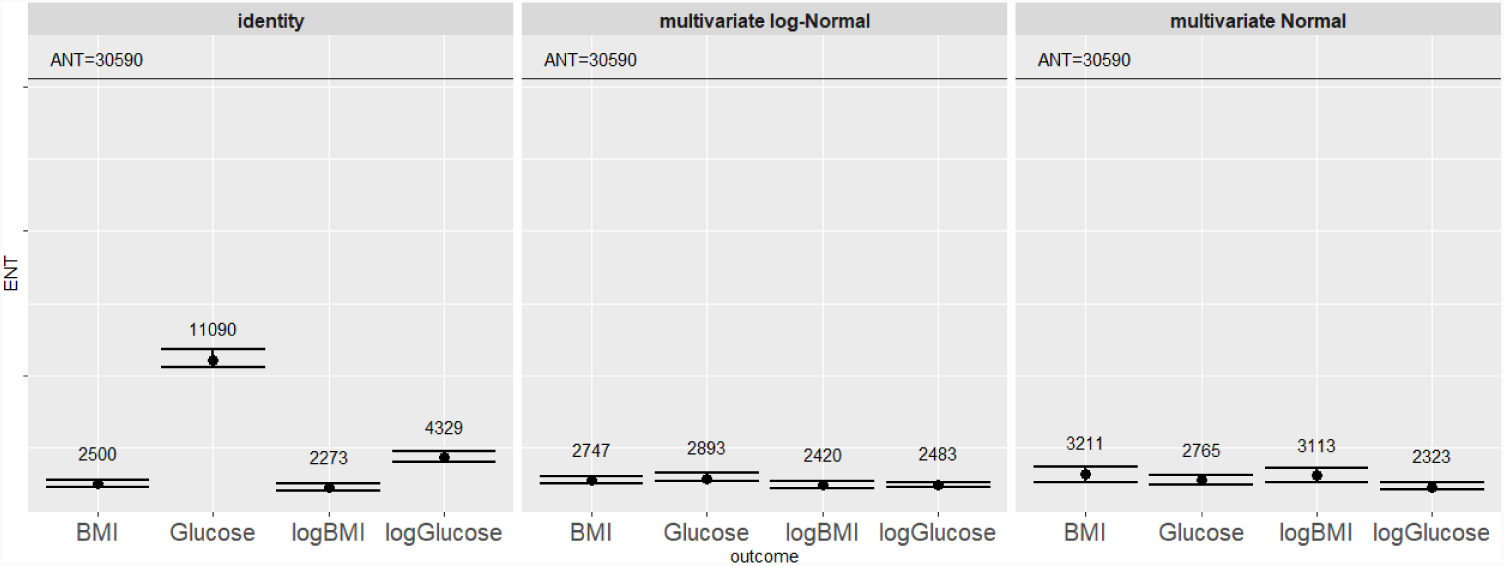
NOESY data: ENT across clinical outcome measures and for different transformations of the features: no transformation (identity), multivariate Normal, multivariate log-Normal. Error bars represent 95% confidence limits. *K*=10,000 permutations.

### 3.2 Simulation study

We now investigate how the the correlation level among features and between outcome and features impact the ENT estimation. To do so we simulate various sets of features with a defined correlation level as follows.

#### 3.2.1 Feature simulation

- Step (1): Generate a square (*M × M*) correlation matrix *A* assuming all variables have unit variance, i.e. the *M* elements on the diagonal are 1s. The [*M* (*M* − 1)]/2 elements of the upper triangular matrix are sampled from Uniform distributions bounded by a certain interval, e.g. high correlation level within the interval [0.75,0.85], medium correlation in [0.45,0.55], or low correlations in [0.25,0.35]. The lower triangle elements are copied from the upper triangle.
- Step (2): As the eigenvalues of *A* are required to be greater than zero, compute *S* as the nearest positive definite matrix to the correlation matrix *A* achieving {min *‖A − S‖*_*F*_: *S* is a correlation matrix}, where 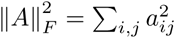 as described by Higham (2002).
- Step (3): Derive the lower triangular matrix *L* via Cholesky decomposition of matrix *S* such that *S* = *LL*’.
- Step (4): *M* multivariate Normal features with zero means result from the product *ZL* between the (*n × M*) matrix *Z* of *M* random N(0,1) i.i.d. features, and the (*M × M*) lower triangular matrix *L*. The correlations of the simulated features are very close to those assigned in matrix *A*.

#### 3.2.2 Outcome simulation

The set of features are analysed together with continuous outcome measures uncorrelated or correlated to the features.

Uncorrelated outcomes of different shapes are easily simulated via parametric distributions such as the Normal distribution for a symmetric outcome, the Skew-Normal distribution for a left skewed outcome, and a Weibull distribution for a right skewed outcome. Figure 2S describes the shapes of the distributions of the simulated uncorrelated outcomes. Figure 4 provide the ENT estimates in the case of correlated features simulated according to Section 3.2.1 and uncorrelated outcomes.

**Figure 4:**
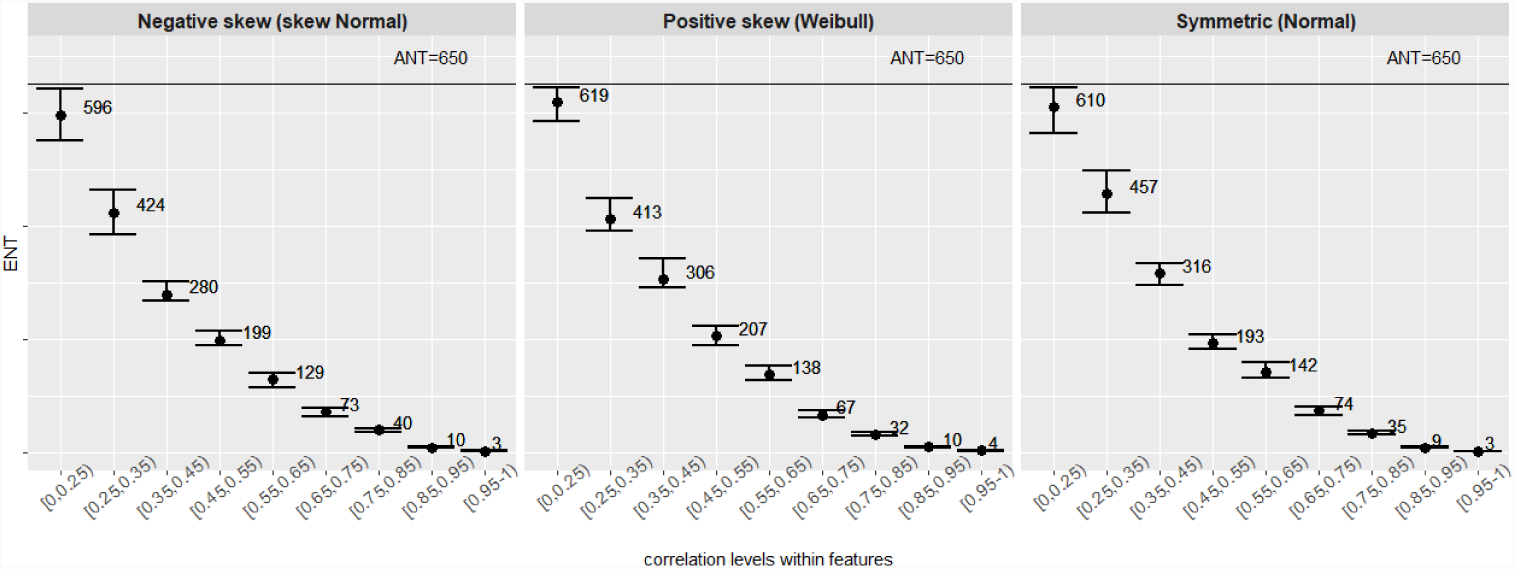
ENT for uncorrelated outcomes across correlated features. Error bars represent 95% confidence limits. *K*=5,000 permutations.

Simulated correlated outcomes can be obtained as a linear combination of few features such as 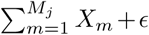, with *ϵ ∼ U* (0, 1) and *M*_*j*_ < *M*, or via procedures based on Cholesky decomposition such as the method used to generate the simulated correlated features described in Section 3.2.1. Figure 3S describes the shapes of the distributions of the simulated correlated outcomes. Figure 5 provides the ENT estimates in the case of correlated simulated features and correlated outcomes.

**Figure 5:**
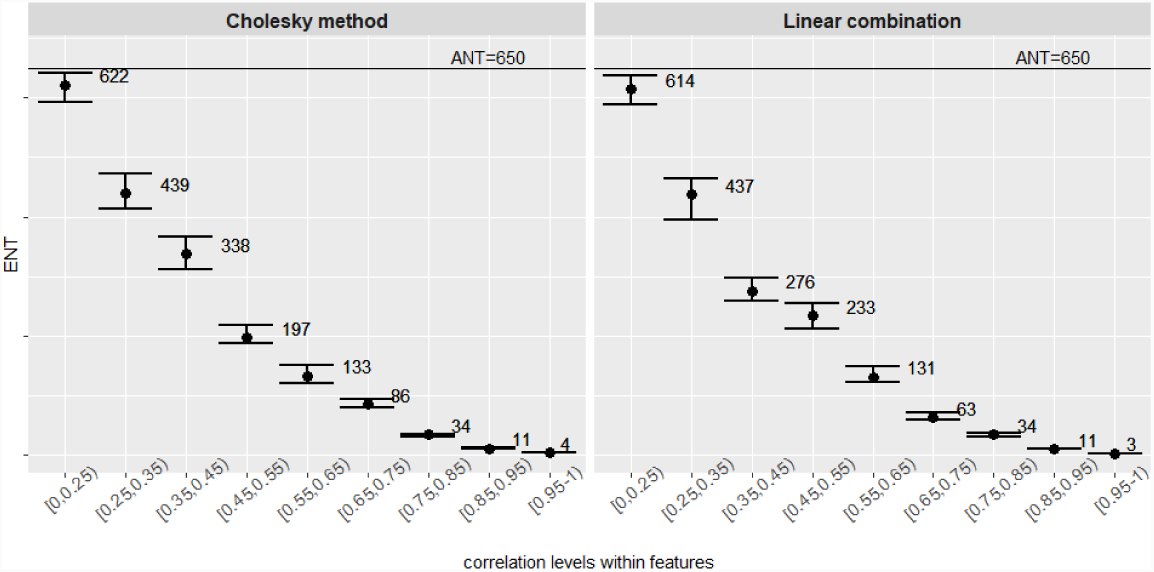
ENT for correlated outcome across correlated features. Error bars represent 95% confidence limits. *K*=5,000 permutations.

We observe that correlation to the outcome makes no discernable difference to relationship between ENT and feature-feature correlation.

The relationships shown in Figure 4 and Figure 5 could be well modelled by polynomial curves (see Supplementary Figure S4 and Supplementary Figure S5).

## 4 Extension and Validation of the approach

A type I error (false-positive) occurs when a true null hypothesis is being rejected. To check whether the procedure accounts for the FWER at the *α* level we measure the the type I error rate within the permutation procedure as the occurrences of having a p-value less or equal than the MWSL.

At this stage we also test the procedure to account for various types of outcomes. In particular, we simulate a total of four uncorrelated outcome measures: a continuous outcome from a Normal distribution, a discrete-binary outcome from a Binomial distribution, a discrete-count outcome from a Poisson distribution, and a time-to-event survival outcome from the Cox proportional hazards model as in Bender *et al*. (2005).

Figure 6 shows the ENT estimates for the BINNED set of features, across a range of simulated uncorrelated outcome measure: continuous, discrete-binary, discrete-count, survival. The procedure effectively controls the FWER at the (default) *α*-level of 5%.

**Figure 6:**
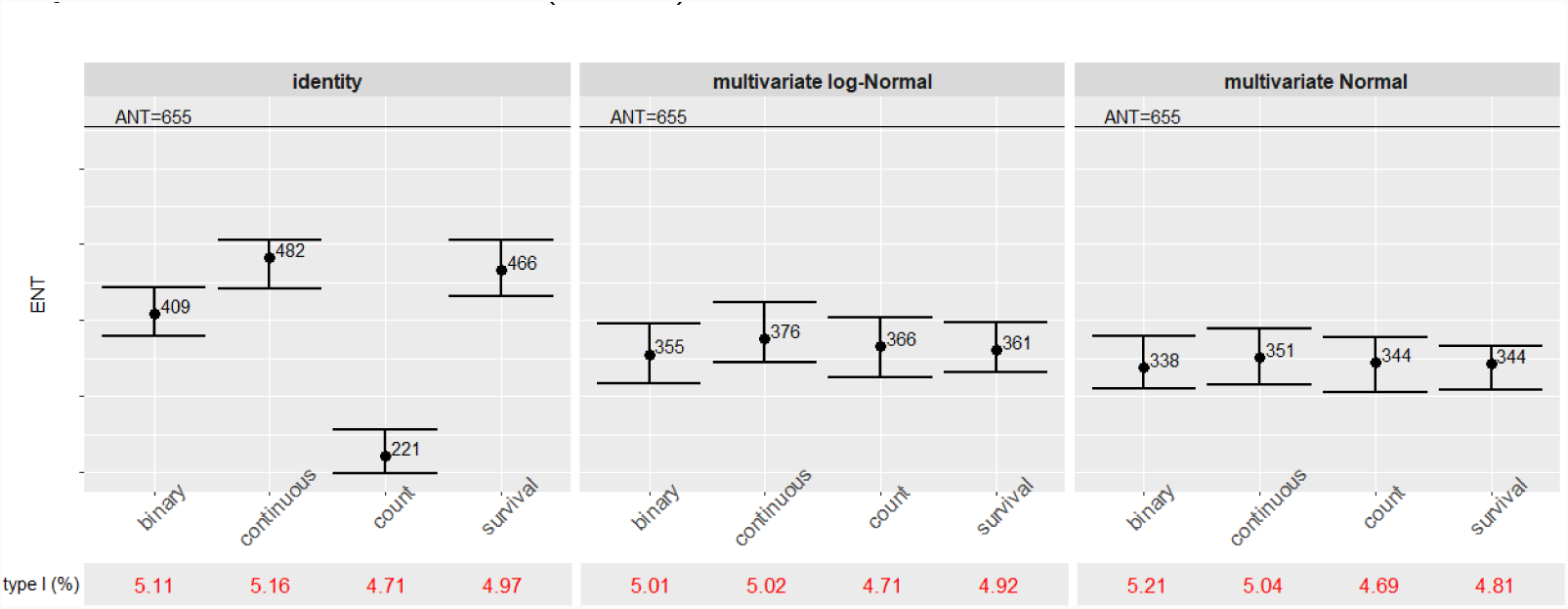
BINNED data: ENT and type I error estimation from the permutation procedure for different types of simulated outcome measures: continuous, discrete-binary, discrete-count, time-to-event survival. *K*=5,000 permutations.

### 4.1 Nonparametric simulation study

To further validate the results we simulate the features based on the original data via a nonparametric approach using PCA. This allowed us to account for the original structure of the data, but does not involve bootstrap or permutation methods (Hastings *et al*., 1999). The algorithm can be described as follows.

- Step (1): By randomly sampling *n*_*t*_ observations from the original data matrix of features *X*, construct the (*n*_*t*_ *× M*) *test* set of features *X*_*t*_, and the 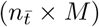 *nontest* set 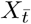, with *n*_*t*_ *< n* and 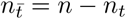.
- Step (2): Standardise the *test* and the *nontest* set by subtracting its respective vector of column means i.e. *µ*_*t*_ and 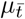, and dividing by its standard deviations i.e. *σ*_*t*_ and 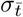, to respectively obtain *Z*_*t*_ and 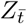.
- Step (3): Compute PCA over the *nontest* set by applying singular value decomposition (SVD) such that 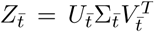, where 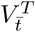 is the (*M × M*) matrix of loadings, while the PC scores are obtained as the product between the 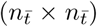 matrix 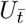 of eigenvectors of 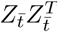, and the 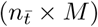 diagonal matrix 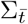.
- Step (4): Use the *nontest* PC scores 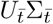 from Step (3) to compute the (*n*_*t*_ *× M*) matrix 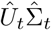 of PC predicted scores for the *test* set.
- Step (5): Build the (*n*_*t*_ *× M*) simulated *test* set of features 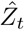 as the product of the predicted scores from Step (4), and the matrix of loadings 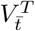 from Step (3) such that 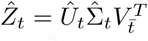. We note that *S* PCs, with *S ≤ M,* can be selected to be used in the predictions, thus 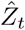 would result from the product of the (*n*_*t*_ *× S*) matrix of PCs and the (*S × M*) matrix of loadings.
- Step (6): From the simulated *test* set of standardised features 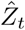 compute the (*M × M*) set of simulated features as 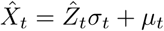.

To simulate features in this context the sample size of the data should be large enough for the data to be split between the test and the nontest set, and no missing values are allowed. A further extension of this method would consider the NIPALS (Nonlinear Iterative Partial Least Squares) algorithm as a modified PCA to accommodate missing values (Martens *et al*., 2001). We consider the BINNED data and we define the *test* and the *nontest* set by randomly sampling *n*_*t*_ = 1, 500 and 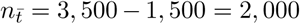 individuals, respectively. From the PCA on the *nontest* set we select 350 PCs to be used to build the simulated *test* set of features 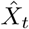 as these account for the 99% of variance explained and are in number close to the average ENT selected by the permutation procedure.

The permutation procedure therefore applied across diverse simulated uncorrelated outcome measures: continuous, discrete-binary, discrete-count and time-to-event survival. Figure 7 confirms that the MWSL procedure effectively controls the FWER at the (default) *α*-level of 5%.

**Figure 7:**
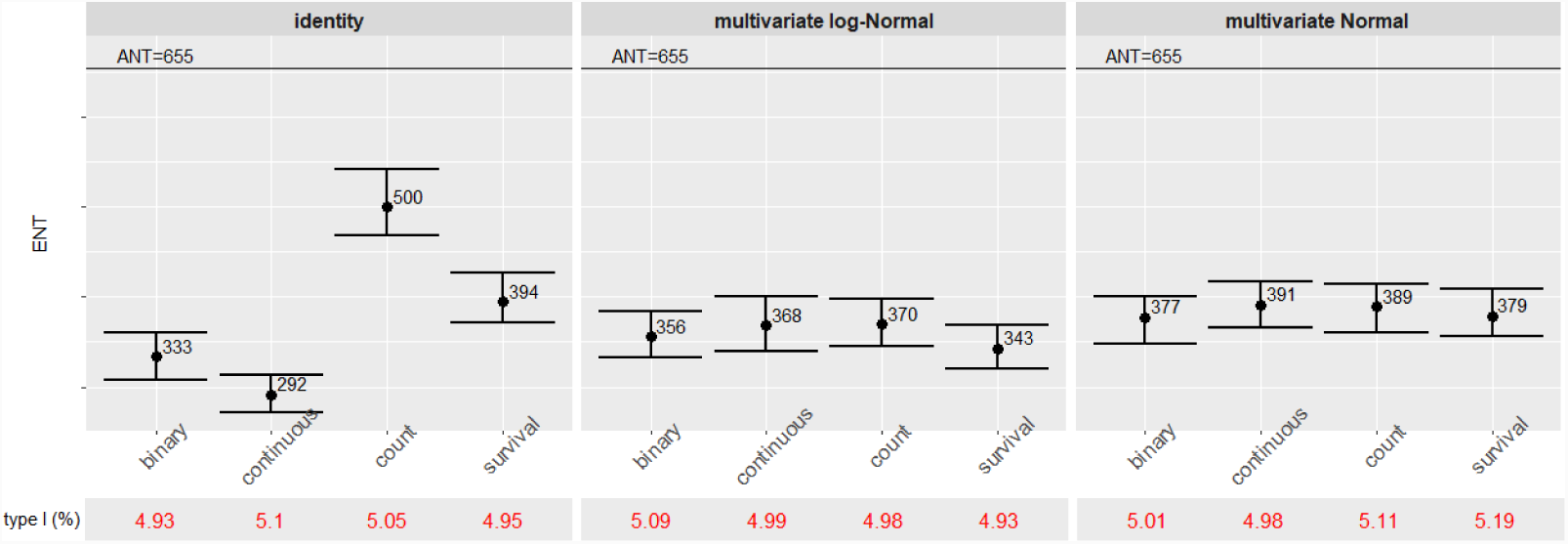
PCA simulated features (M=655, *n*_*t*_=1,500, PCs=350). ENT estimates with 95% confidence intervals and type I error estimation from the permutation procedure for different type of simulated outcome measures: continuous, discrete-binary, discrete-count, time-to-event survival. *K*=5,000 permutations.

## 5 Conclusions

In this paper we consider the empirical estimation of a significance level threshold for multiple testing adjustment when dealing with correlated data, as commonly found in omics studies.

The procedure is based on an iterative permutation approach via univariate regression models. The features are simulated via parametric methods such as multivariate Normal and multivariate log-Normal distributions to retain the dependence structure in the data. When the permutation procedure is applied to the approximated data the MWSL is stable across outcome measures with diverse properties.

Spectral variables from metabolic profiles exhibit a high degree of collinearity, and this is supported by our finding that in all scenarios considered, when parametric methods are applied to approximate the structure of the data, the MWSL estimated through the permutation procedure is larger than the threshold obtained via a metabolome-wide Bonferroni or Sidak corrections. Therefore, the corresponding ENT is always less than the actual number of tests as it mainly depends on the extent of correlation within the data.

The extent of collinearity is summarized by the *R* ratio of effective to actual number of tests. For the examples in this paper, *R* was found to be 55% for the CPMG data (high-resolution and and BINNED version), and around 10% for the NOESY resolution. This is consistent with the expected higher degree of correlations between spectral variables in the NOESY data.

The number of independent variables might give an indication of the number of independent observable metabolic processes exhibited by the system, since each independent process might be expected to manifest itself through multiple metabolic variables. If the data are interpreted this way, our analysis suggests that around 350 separate metabolic processes for the BINNED resolution, around 15,000 processes within the CPMG resolution and 3,000 processes for the NOESY resolution are being captured by NMR metabolic profiling of human serum.

Via simulation and real data study it was shown that the permutation procedure performs satisfactorily in the task of discovering associations, while controlling the false positive rate at the desired level.

## Availability

The `MWSL` R package is available at https://github.com/AlinaPeluso/MWSL

Within the package the lower resolution CPMG data referred to in the text as BINNED data is also available.

## Supplementary material

Details about the data used in the analysis in the form of figures and tables are available as supplementary information.

## Funding and acknowledgements

We thank Dr Marc Chadeau-Hyam and Dr Raphaele Castagne for useful discussions. This work was undertaken as part of the PhenoMeNal project (Horizon2020, 2015-2018), European Commission grant EC654241.

## References

Holmes E. (2008) Human metabolic phenotype diversity and its association with diet and blood pressure, Nature, 453, 396–400.

Hoggart, Clive J. and Clark, Taane G. and De Iorio, Maria and Whittaker, John C. and Balding, David J. (2008) Genome-wide significance for dense SNP and resequencing data, Genetic Epidemiology: The Official Publication of the International Genetic Epidemiology Society, 32 (2), 179–185.

Chadeau-Hyam, Marc and Ebbels, Timothy M.D. and Brown, Ian J. and Chan, Queenie and Stamler, Jeremiah and Huang Chiang Ching and Daviglus, Martha L. and Ueshima, Hirotsugu and Zhao, Liancheng and Holmes, Elaine and others (2010) Metabolic profiling and the metabolome-wide association study: significance level for biomarker identification, Journal of Proteome research, 9 (9), 4620–4627.

Castagnée, Raphaële and Boulangé Claire Laurence and Karaman, Ibrahim and Campanella, Gianluca and Santos Ferreira, Diana L. and Kaluarachchi, Manuja R. and Lehne, Benjamin and Moayyeri, Alireza and Lewis, Matthew R. and Spagou, Konstantina and others (2017) Improving Visualization and Interpretation of Metabolome-Wide Association Studies: An Application in a Population-Based Cohort Using Untargeted 1H-NMR Metabolic Profiling, Journal of Proteome research, 16 (10), 3623–3633.

Paparoditis, Efstathios and Politis, Dimitris N. (1999) The local bootstrap for periodogram statistics, Journal of Time Series Analysis, 20 (2), 193–222.

Schäafer, Juliane and Strimmer, Korbinian (2005) A shrinkage approach to large-scale covariance matrix estimation and implications for functional genomics, Statistical applications in Genetics and Molecular Biology, 4 (1).

Bild, D. E. and Bluemke, D. A. and Burke, G. L. and Detrano, R. and Diez Roux, A. V. and Folsom, A. R. and Greenland, P. and Jacob, D. R. and Kronmal, R. and Liu, K. (2002) Multi-ethnic study of atherosclerosis: objectives and design, Am. J. Epidemiol., 156 (9), 871–881.

Karaman, I. and Ferreira, D. L. S. and Boulangé, C. L. and Kaluarachchi, M. R. and Herrington, D. and Dona, A. C. and Castagné, R. and Moayyeri, A. and Lehne, B. and Loh, M. (2016) Workflow for Integrated Processing of Multicohort Untargeted (1)H NMR Metabolomics Data in Large-Scale Metabolic Epidemiology, J. Proteome Res., 15 (12), 4188–4194.

Cheverud J. M. (2001) A simple correction for multiple comparisons in interval mapping genome scans, Heredity, 87, 52–58.

Nyholt D. R. (2004) A simple correction for multiple testing for single-nucleotide polymorphisms in linkage disequilibrium with each other, Am. J. Hum. Genet., 74, 765769.

Li J., and Ji L. (2005) Adjusting multiple testing in multilocus analyses using the eigenvalues of a correlation matrix, Heredity, 95, 221–227.

Higham, Nicholas J. (2002) Computing the nearest correlation matrix - a problem from finance, Journal Name, 22 (3), 329–343.

Bender R., and Augustin T., and Blettner, M. (2005) Generating survival times to simulate Cox proportional hazards models, Statistics in medicine, 24(11), 1713–1723.

Hastings W. K. (1999) Monte Carlo sampling methods using Markov chains and their applications, Biometrika, 57, 97–109.

Martens H., and Martens M. (2001) Multivariate analysis of quality: an introduction, John Wiley & Sons, 381.

PhenoMeNal (Phenome and Metabolome aNalysis): Large-scale Computing for Medical Metabolomics (2015-2018) European Commission’s Horizon2020, https://phenomenal-h2020.eu/.

