## Supplementary material for "Estimation of permutation-based metabolome-wide significance thresholds"

January 13, 2019

[1] Faculty of Medicine, Department of Surgery & Cancer, Imperial College London, South Kensington Campus, SW7 2AZ, London, UK.
[2] Centre for Molecular Informatics, Department of Chemistry, University of Cambridge, Lensfield Road, CB2 1EW, Cambridge, UK.
*To whom correspondence should be addressed:
{a.peluso, r.glen, t.ebbels}@imperial.ac.uk.

Additional information are available at https://github.com/AlinaPeluso/MWSL.

### List of Tables

1 Descriptive statistics for the clinical outcome measures. . . . . . . . . . . . . . . . . . . . 2
2 Descriptive statistics for the fixed effects covariates. . . . . . . . . . . . . . . . . . . . . . 2

The outcomes of interest are glucose concentrations and the body mass index (BMI). Table S1 presents the descriptive statistics for the clinical outcome measures in Figure S1, while Table S2 reports the descriptive statistics for the fixed effects covariates used in the study.

Table 1: Descriptive statistics for the clinical outcome measures.

| | mean | sd | median | min | max | skewness | kurtosis |
|---|---|---|---|---|---|---|---|
| Glucose (mg/dL) | 98.28 | 31.10 | 90 | 38 | 507 | 4.17 | 28.89 |
| Logarithm of Glucose | 4.56 | 0.23 | 4.5 | 3.64 | 6.23 | 2.22 | 10.35 |
| BMI (kg/m$^2$) | 28.14 | 5.39 | 27.34 | 15.36 | 61.86 | 46.50 | 4.45 |
| Logarithm of BMI | 3.32 | 0.18 | 3.31 | 2.73 | 4.12 | 1.39 | 3.20 |

Table 2: Descriptive statistics for the fixed effects covariates.

| | mean | sd |
|---|---|---|
| Age (years) | 62.89 | 10.32 |
| Gender | 0.51 | 0.49 |
| Height (cm) | 166.43 | 10.23 |
| Ethnicity: Caucasian | 0.39 | 0.49 |
| Ethnicity: Hispanic | 0.23 | 0.42 |
| Ethnicity: African-American | 0.25 | 0.43 |
| Ethnicity: Chinese-American | 0.13 | 0.34 |
| Smoking: Never | 0.51 | 0.50 |
| Smoking: Former | 0.12 | 0.33 |
| Smoking: Current | 0.38 | 0.48 |
| LDL cholesterol (mg/dL) | 117.67 | 31.04 |
| HDL cholesterol (mg/dL) | 51.29 | 14.41 |
| Systolic blood pressure (mmHg) | 126.92 | 21.54 |
| Blood pressure treatment | 0.38 | 0.49 |
| Diabetes | 0.14 | 0.34 |
| Lipids treatment | 0.17 | 0.37 |

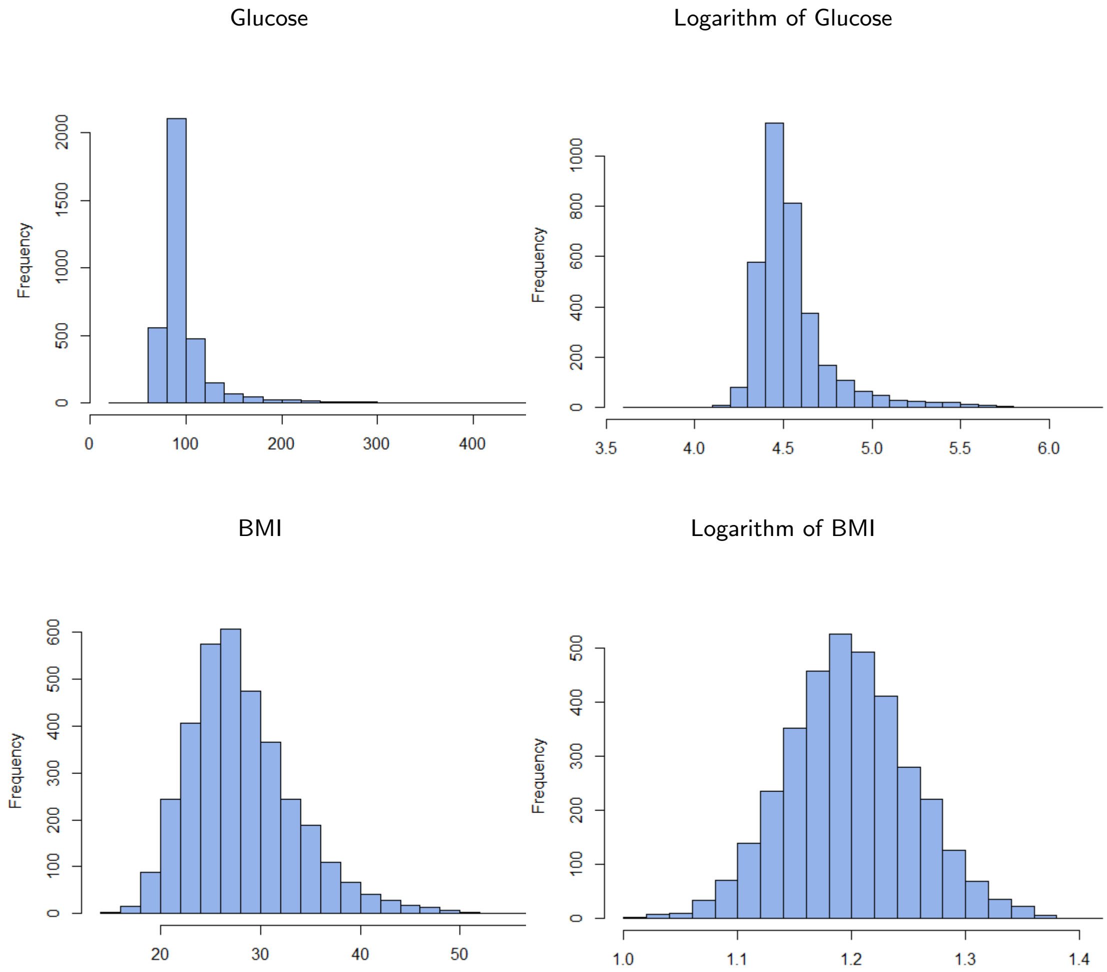

Figure 1: Distributions of the clinical outcome measures.

Uncorrelated outcomes of different shapes are easily simulated via parametric distributions such as the Normal distribution for a symmetric outcome, the Skew-Normal distribution for a left skewed outcome, and a Weibull distribution for a right skewed outcome. Figure S2 describes the shapes of the distributions of the simulated uncorrelated outcomes.

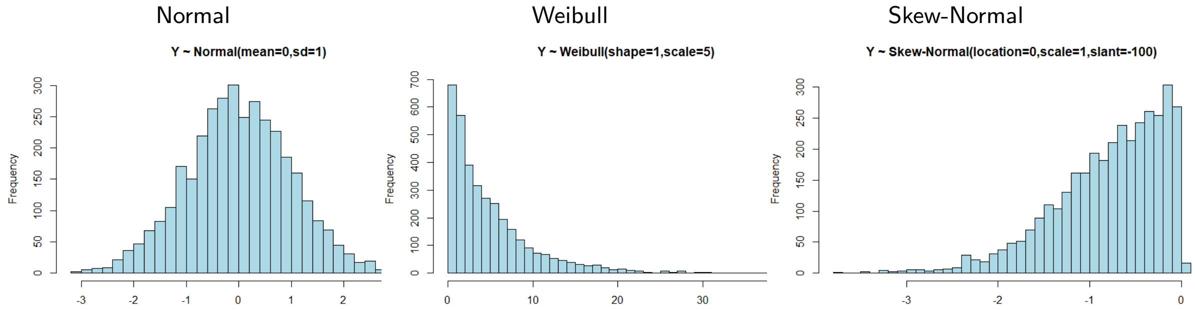

Figure 2: Distributions of the simulated uncorrelated outcome measures.

Figure S3 describes the shapes of the distributions of the simulated correlated outcomes obtained as a linear combination of few features such as $\sum_{m=1}^{M_j} X_m + \epsilon$, with $\epsilon \sim U(0,1)$ and $M_j < M$, or via procedures based on Cholesky decomposition.

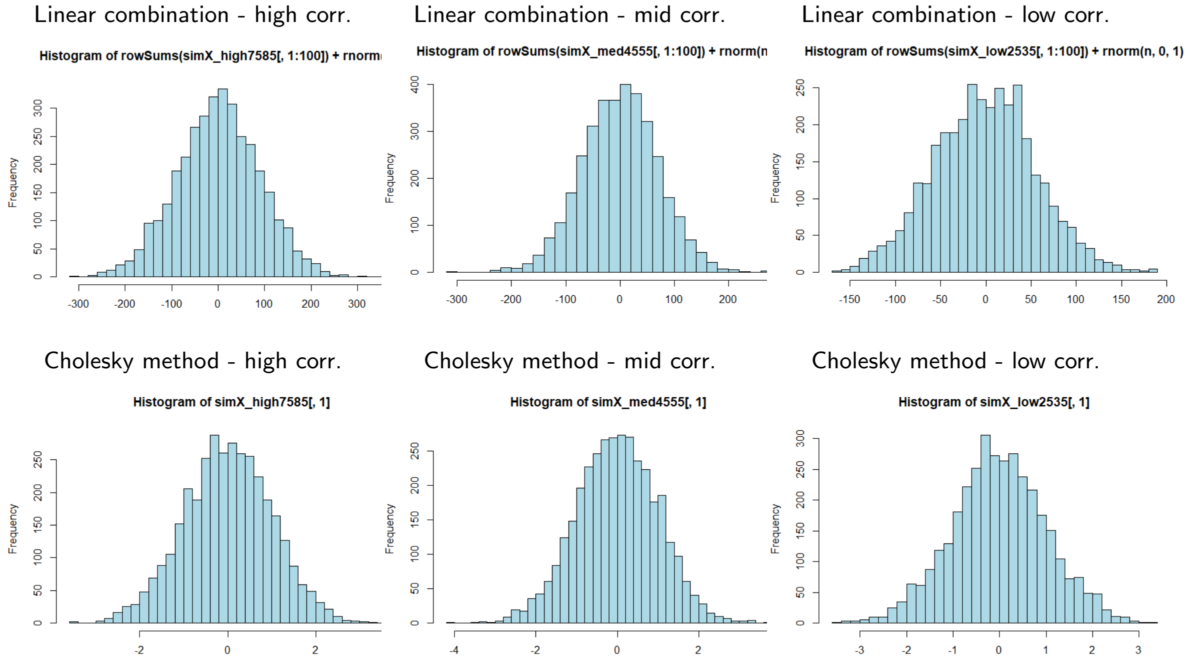

Figure 3: Distributions of the simulated correlated outcome measures.

To describe the inverse relationship between the correlation levels of the features and the ENT estimates i.e. decreasing R(%) ratios for higher level of correlations within the features, we fit a nonlinear 3rd degree-polynomial curve to the ENT estimates as showed in Figure S4 and Figure S5 in the case of correlated features and correlated and uncorrelated outcomes, respectively.

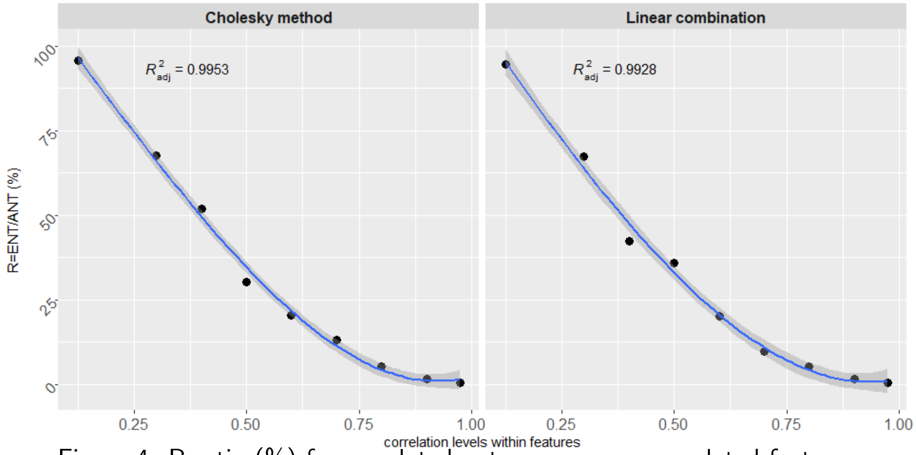

Figure 4: R ratio (%) for correlated outcomes across correlated features.

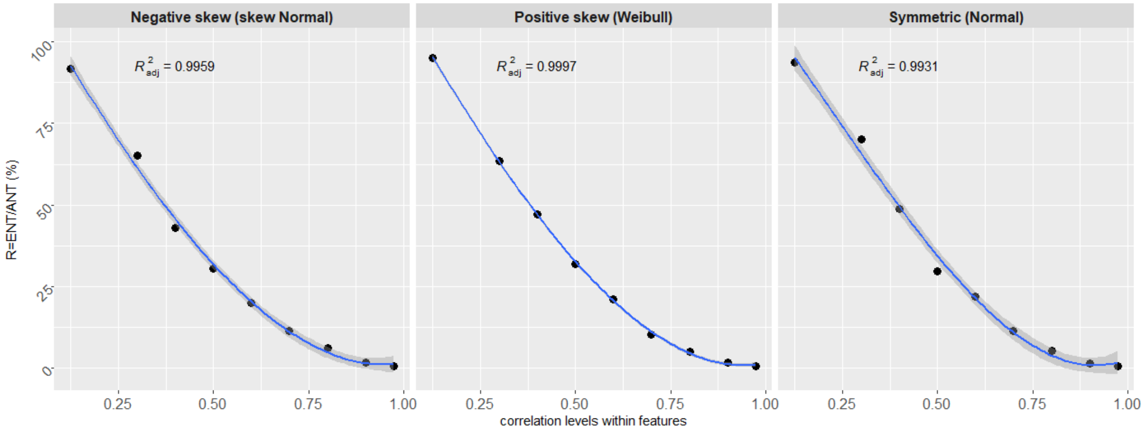

Figure 5: R ratio (%) for uncorrelated outcomes across correlated features.
